# Tough Hydrogel-Based Biocontainment of Engineered Organisms for Continuous, Self-Powered Sensing and Computation

**DOI:** 10.1101/2020.02.11.941120

**Authors:** Tzu-Chieh Tang, Eleonore Tham, Xinyue Liu, Kevin Yehl, Alexis J. Rovner, Hyunwoo Yuk, Farren J. Isaacs, Xuanhe Zhao, Timothy K. Lu

## Abstract

Genetically modified microorganisms (GMMs) can enable a wide range of important applications, including environmental sensing, precision therapeutics, and responsive materials. However, containment of GMMs to prevent environmental escape and satisfy regulatory requirements is a bottleneck for real-world use^1–7^. While biochemical strategies have been developed to restrict unwanted growth and replication of GMMs in the environment^8–12^, there is a need for deployable physical containment technologies to achieve redundant, multi-layered, and robust containment^2^. In addition, form factors that enable easy retrieval would be useful for environmental sensing. To address this challenge, we developed a hydrogel-based encapsulation system for GMMs that incorporates a biocompatible multilayer tough shell and an alginate-based core. This DEployable Physical COntainment Strategy (DEPCOS) allows no detectable GMM escape, bacteria to be protected against environmental insults including antibiotics and low pH, controllable lifespan, and easy retrieval of genetically recoded bacteria. To highlight the versatility of a DEPCOS, we demonstrate that robustly encapsulated cells can execute useful functions, including performing cell-cell communication with other encapsulated bacteria and sensing heavy metals in water samples from the Charles River. We envision that our multilayered physical and chemical containment strategy will facilitate the realization of a wide range of real-world applications for ‘living’ biosensors.

---

Genetically modified microorganisms (GMMs) are being developed and used for bioremediation^13^, agriculture^14^, and the production of biofuels^15^ and pharmaceuticals^16^. However, the potential for GMMs to escape into the environment has created a need for strategies to contain these organisms and prevent their uncontrolled release.

Chemical biocontainment utilizes chemical barriers to impede the escape and survival of microorganisms in the environment^4–6^. Several strategies have been developed for chemical containment of GMMs. For example, GMMs can carry gene circuits that require specific chemical combinations to prevent cell death by inhibiting toxin production^10^, rescue cells from being killed by a constitutively expressed toxin by producing the corresponding antitoxin^17^, or multi-layered safeguards that modulate the expression of essential genes and toxins^18^. In addition, microbes can be engineered with auxotrophies so that they require synthetic amino acids for survival^19–21^. However, chemical strategies alone are imperfect for containment because mutation rates of GMMs, while low, are never zero, thus resulting in escape mutants. This implies that the number of chemically-contained GMMs that can be deployed is intrinsically limited by its mutation rate^21^. Thus, it would be beneficial to combine biological containment strategies and physical encapsulation such that functional redundancy further reduces any chance of inadvertent escape.

To address this challenge, we created a DEployable Physical COntainment Strategy (DEPCOS) that prevents GMM escape while providing a tunable protective environment in which GMMs can execute engineered functions. DEPCOS erects a physical barrier to prevent GMMs from escaping into their surroundings, limit horizontal gene transfer between GMMs and natural species in the environment, and allow for easy retrieval of bacterial communities.

Hydrogels are desirable materials for encapsulating living cells as they provide an aqueous environment that can be infused with nutrients, allowing for cell growth^22,23^ and sensing^24^, while also protecting against environmental hazards^25^. Alginate forms hydrogels in the presence of di-cationic solutions (e.g., Ca^2+^, Ba^2+^) and has been used in various biomedical applications^26–30^ because of its low cost, negligible cytotoxicity, and mild gelation conditions. However, weak mechanical properties and susceptibility to multiple chemical conditions (such as low pH, citrate, and phosphate) make alginate, as well as other traditional hydrogels, poor solutions for robust physical containment when used on their own^31–33^.

Core-shell designs that include an alginate core and a polymer-based protective shell have emerged as potential design for alginate-based microbial biosensors^32,34^. Nonetheless, there is a major need for a mechanically-tough shell that is also highly permeable to analytes for sensing. Our DEPCOS design for bacterial encapsulation consists of two parts: 1) an alginate-based hydrogel core and 2) a tough hydrogel shell (Fig. 1a) that combines both a stretchy polymer network (polyacrylamide) and an energy dissipation network (alginate, through the unzipping of ionic crosslinking between polymer chains)^35^. This shell material is extremely tough and resistant to fracture, yet retains permeability for small molecules^36^. Herein, we test the biocompatibility of this hydrogel and further expand upon its physical characterization for core-shell particle form factors.

**Fig. 1.**
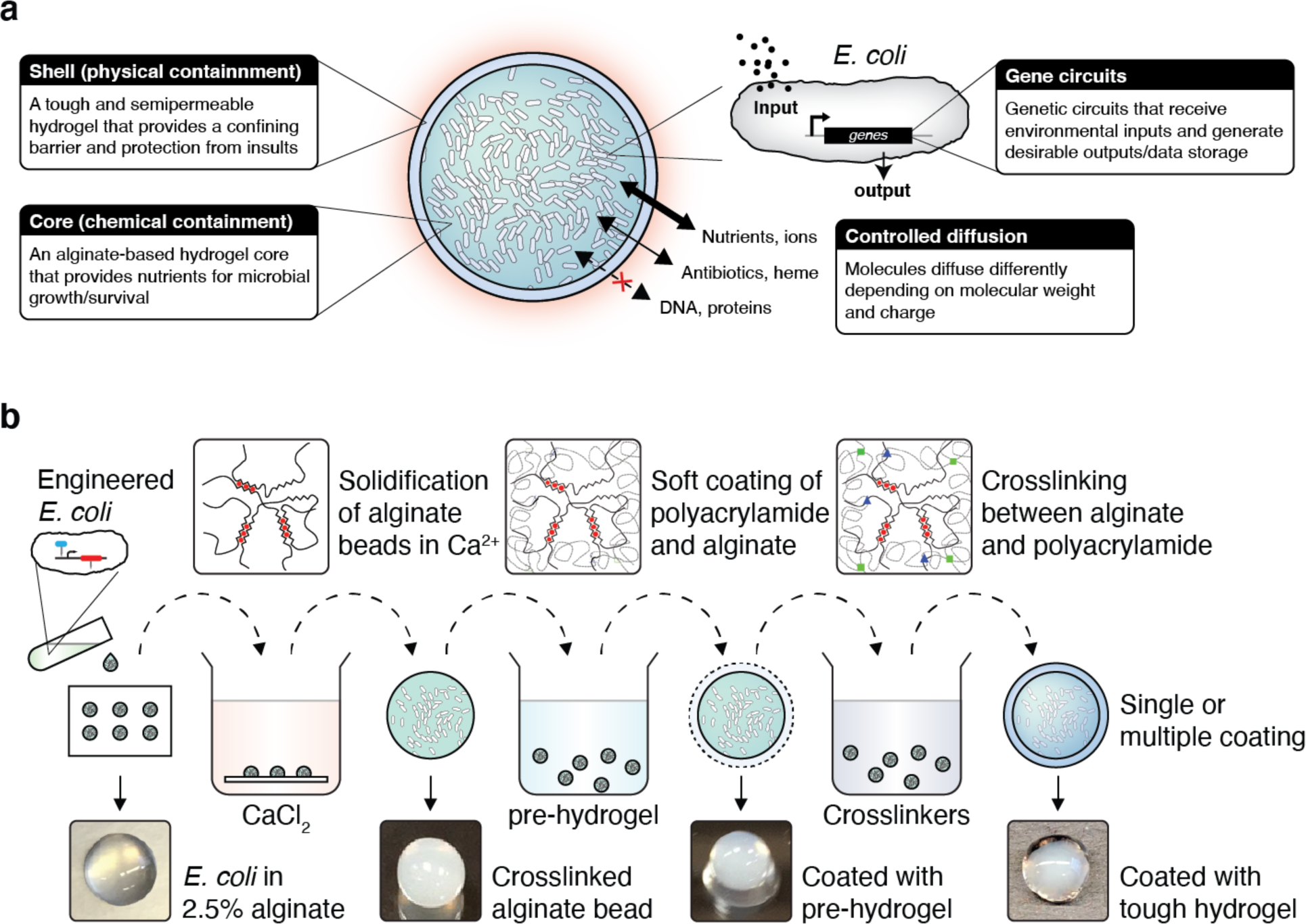
Cell encapsulation in tough hydrogel capsules. **(a)** Schematic of the DEPCOS platform. **(b)** The process of coreshell encapsulation of cells. Droplets of 2.5% alginate with engineered *E coli* were crosslinked in a calcium solution to form the soft core of the beads, which were then coated with a layer of alginate/polyacrylamide to form a tough hydrogel shell. The process can be repeated to achieve multiple coatings.

To incorporate living cells into the particle core, a liquid culture of *E. coli* was mixed with alginate in 50 μl or 100 μl droplets that were crosslinked with calcium ions to form spheres. The cell-containing alginate hydrogel was easily shaped by a mold or cut into different geometries (Supplementary Fig. 1). Cores were then coated with the tough polyacrylamide-alginate hydrogel layer^35^ (Fig. 1b). In our core-shell system, the alginate core is pre-loaded with nutrients to support growth while the hydrogel shell provides mechanical protection for the entire bead. For downstream analyses after deployment, cells can easily be retrieved from the beads by removing the shell with a razor blade and homogenizing the core (Supplementary Fig. 2).

We hypothesized that the tough hydrogel layer would serve as a containment mechanism because its pore size (5-50 nm) is too small for *E. coli* to penetrate^37,38^. To test this hypothesis, we measured the containment efficiency of hydrogel beads by incubating *E coli*-encapsulated beads at the optimal temperature for bacterial growth (37°C) with shaking. Specifically, we encapsulated a concentration of ~10^9^ bacteria/mL in each bead. Beads lacking a tough shell allowed bacteria to escape into the surrounding media and to grow to high densities after overnight incubation, whereas there was no physical escape of bacteria from coated beads even after 72 hours of incubation (Fig. 2a and Supplementary Fig. 3). In this assay, we plated all of the media (5 mL) surrounding the beads, with a lower limit of detection (LLOD) of 1 CFU in 5 mL.

**Fig. 2.**
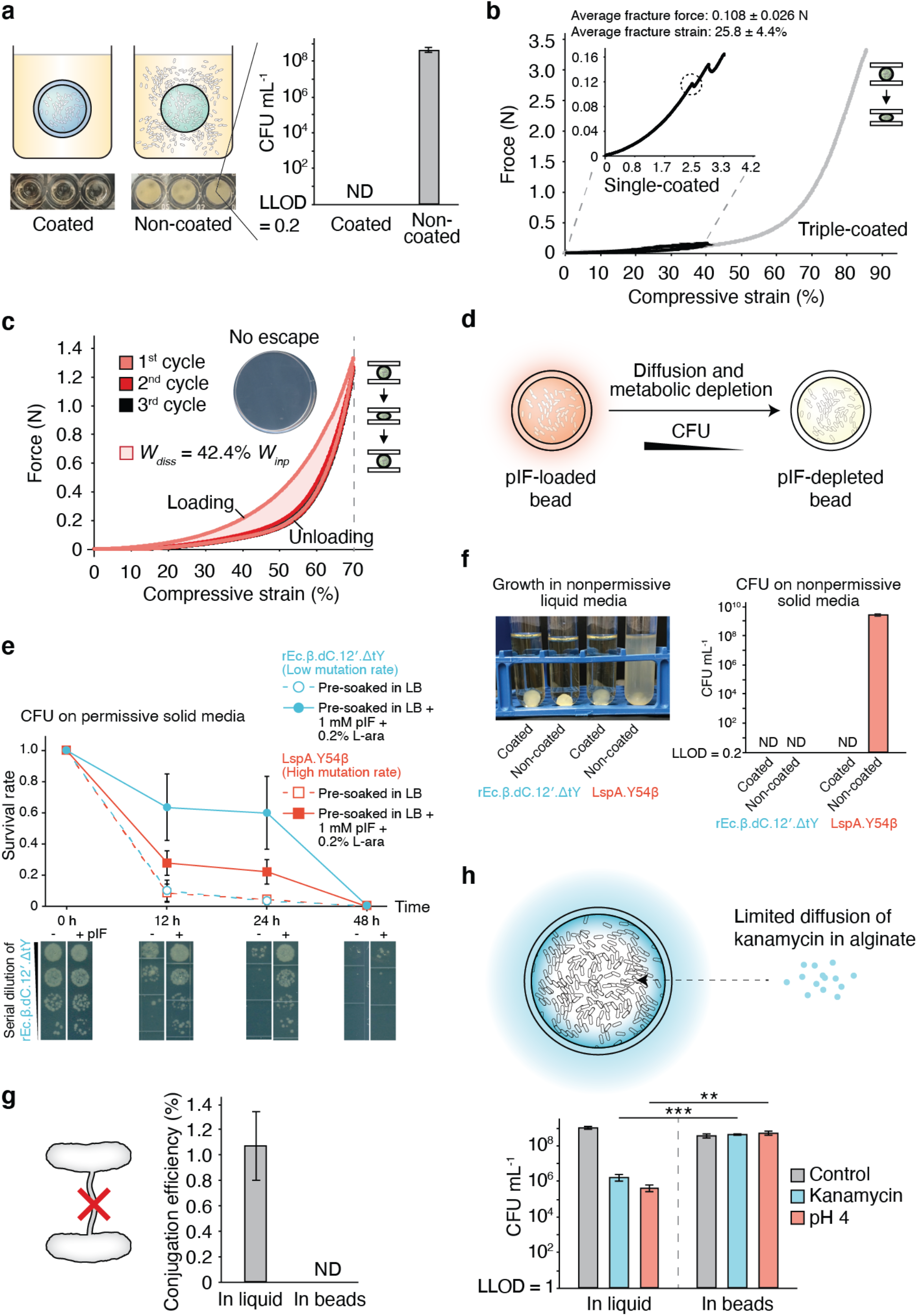
Combining chemical and physical strategies for optimal biocontainment. **(a)** Encapsulated bacteria escaped from non-coated beads at high rates but did not escape from tough-hydrogel-coated beads at detectable levels after 72h. Inset shows media in which non-coated and coated bead were grown for 24h (lower limit of detection (LLOD) = 1 CFU / 5 mL). ND = not detectable. **(b)** Typical force-displacement curves of single-layer tough-hydrogel-coated beads subjected to 40% (black) compressive strain and triple-coated beads subjected to compression up to 85% (gray) compression. Inset zooms in on the single-coated bead stress-strain curve. The average maximum strain and force before fracture for the single-layer coating were 25.8 % and 0.108 N, respectively. Triple-coated beads showed no fracture under compression. **(c)** Cyclic compression of triple-layer coated beads showed hysteresis in the stress-strain curve between the first and second cycles due to plastic deformation. Energy dissipated E_diss_ in the first cycle was calculated as 42.4% of the total work E_in_. Stress-strain curves are representative of at least 6 independent experiments. Inset shows there was no escape that plating the surrounding media of a bead after cyclic compression yielded no colonies. **(d)** Schematic of combining chemical and physical biocontainment strategies. Number of viable cells in the beads decreases as pIF in permissive media is consumed or diffuses away. **(e)** Comparison of cell survival in beads between two GRO strains with different containment efficiencies. Survival rate calculated by the number of encapsulated GRO in tough-hydrogel beads that were pre-soaked in permissive media (LB + 1 mM pIF + 0.2% L-ara) (closed circles and solid lines) versus encapsulated GRO in tough-hydrogel beads that were pre-soaked in nonpermissive media (LB only) (open circles and dashed lines) before incubation of the beads in LB. Survival rates were calculated by normalizing colony forming units (CFU) from samples inside the beads, plated on permissive solid media at each time point to CFU at 0 h. Dilution series of the rEc.*β*.dC.12’. ΔtY at different incubation time points are shown at the bottom. Data are mean ± s.d. (n ≥ 3). **(f)** Left: Escape of GROs into 5 mL of nonpermissive media (LB only) surrounding the coated versus non-coated beads containing the GROs after shaking the tough-hydrogel beads at 200 rpm for 3 days. Right: The surrounding media was plated on nonpermissive solid media in order to obtain CFU counts. (ND: not detectable with LLOD = 1 CFU/5 mL) Data are mean ± s.d. (n ≥ 5). **(g)** The tough hydrogel shell prevents horizontal gene transfer by direct cell-to-cell conjugation. Conjugation efficiency is calculated as the ratio of recipient strain that acquired the F’ plasmid over the total number of recipient cells in media. ND = not detectable. Data are mean ± s.d. (n ≥ 3). **(h)** Survival of bacteria after subjecting liquid bacterial cultures or bacteria in tough-hydrogel-coated beads to environmental challenges such as antibiotics (30 μg/ml kanamycin for 2 hours), low pH (pH 4 for 4 hours), and untreated controls (LLOD = 200 CFU/mL). Data are mean ± s.d. (n ≥ 3, **p < 0.01, ***p < 0.001; Student’s t-test).

Next, we used compression testing to characterize the mechanical robustness of the hydrogel-bacteria beads (radius = 3-4 mm) with varying shell layers. We found that beads with a single-layer shell coating (Supplementary Fig. 4) could sustain 25% compressive strains and forces up to ~0.1 N before fracture occurred (Fig. 2b and Supplementary Fig. 5a). We further improved the mechanical properties of the beads by creating multilayer shells via repetitive coating. With beads that were coated with three layers of tough polyacrylamide-alginate hydrogel, we did not observe any fracture when the beads were subjected to up to 85% compressive strains and forces up to ~3.3 N (Fig. 2b and Supplementary Fig. 5a). The bead capsules were also subjected to cyclic compression at 70% strain, revealing a pronounced hysteresis due to plastic deformation and energy dissipation (Fig. 2c and Supplementary Fig. 5b). Based on the dimensions of the beads, the cyclic effective compressive stress was calculated as ~70 kPa (Supplementary Fig. 5b), which is equivalent to pressure at ~7 m depth under water and ~4 m depth under dry soil^39^, and is comparable to our previously published ingestible hydrogel device^40^. This result suggests that these beads could sustain much stronger stresses higher than the maximum gastric pressure (~10 kPa)^41^, highlighting their potential for *in vivo* biosensing. Thus, multilayer coating with elastic tough hydrogel around an alginate core provides mechanical robustness to the entire capsule, a phenomenon which is observed with other stiff polymer coatings^42^. Importantly, zero CFU counts were detected when plating the surrounding media that was incubated with compressed beads, suggesting that the capsules withstood successive compressions without fracturing and maintained perfect containment needed for safe environmental deployment (Fig. 2c, inset).

Since extreme forces could potentially compromise our hydrogels and permit bacterial escape, we hypothesized that chemical containment could be employed to enforce an additional layer of control over encapsulated cells. Genomically recoded organisms (GROs, microbes with synthetic autotrophies)^21^ can be contained because the growth of these microbes is dependent on the supply of synthetic amino acids (enabling a permissive environment). Here, we sought to combine physical and chemical strategies for biocontainment by encapsulating two GROs auxotrophic for the synthetic amino acid p-iodo-phenylalanine (pIF) in tough hydrogel beads (Fig. 2d). The *E coli* strains rEc.β.dC.12’.ΔtY (mutation rate <4.9 × 10^−12^) and LspA.Y54β (mutation rate = 1.86 × 10^−5^) have amber codons (TAG) inserted in three (Lsp, DnaX, SecY) and in one (Lsp) essential genes^21^, respectively, to restrict growth to permissive media (containing pIF). We showed that: 1) chemical containment in the beads enables programmable loss of cellular viability after 48 hours, which prevents undesirable growth once a given time frame has expired; and 2) physical containment adds another layer of protection over chemically contained microbes, which is necessary for applications that require extremely high standards of biocontainment.

First, beads encapsulating the pIF-auxotroph GROs, rEc.β.dC.12’.ΔtY and LspA.Y54β (~10^5^ cells per bead), were pre-soaked in LB in the absence (non-permissive media) or presence (permissive media) of 1 mM pIF and 0.2% L-arabinose (L-ara), which is required for aminoacyl-tRNA synthetases (aaRS) expression in these strains^21^. We hypothesized that in non-permissive media, the GROs would be unable to synthesize functional essential proteins and thus, lose viability. Indeed, beads pre-soaked in non-permissive media failed to sustain cell growth and showed less than 10% survival after 12 hours in LB only (Fig. 2e, cells were plated on permissive solid media), with no survival detected at 24 h. On the other hand, pre-soaking encapsulated beads in permissive media greatly prolonged cell survival. Greater than 50% of the rEc.β.dC.12’.ΔtY population and >25% of the LspA.Y54β population remained viable after 24h of incubation. Nearly all cells (>99%) lost viability after 2 days of incubation, which we believe is due to pIF and L-ara depletion by cells, as well as passive diffusion of these molecules out of the encapsulated hydrogel. Because many chemical induction and sensing responses in *E coli* require less than 24 hours to complete, this defined survival window could be used to prevent the undesirable growth of cells upon completion of tasks.

We demonstrated the benefit of combining physical and chemical containment (Fig. 2f) by placing beads in non-permissive liquid media and then plating samples from the liquid media on non-permissive solid media. This experiment allowed us to screen for escape mutants. For beads with tough hydrogel coating encapsulating either GRO strain (1.2 × 10^7^ cells), no viable cells were observed in the surrounding non-permissive media at the end of a 3-day shaking incubation period, indicating complete containment (Fig. 2f). When rEc.β.dC.12’.ΔtY cells (low mutational escape rate, <4.9 × 10^−12^)^21^ were encapsulated in beads without the tough hydrogel coating, no viable cells were observed in the non-permissive media. On the other hand, when LspA.Y54β cells (higher mutational escape rate, ~ 1.86 × 10^−5^)^21^ were encapsulated in beads without the tough hydrogel coating, mutational escape was observed and cells grew in the non-permissive media. These results demonstrate that physical containment can complement chemical containment strategies to achieve near-zero escape rates (chemical plus physical). Furthermore, we can program a “biological timer” system that eliminates potential bacterial growth outside the bead even when the hydrogel shell is compromised.

Horizontal gene transfer (HGT) of engineered genes into the environment and disruption of native ecosystems is a major regulatory concern regarding deployment of GMMs. Since DNA is much smaller than bacteria, we sought to explore whether DEPCOs could prevent HGT. We used a bacterial conjugation assay (Fig. 2g) where the conjugation efficiency of an F-plasmid carrying a chloramphenicol (Cm) resistance from an F+ donor strain was measured for transfer into a recipient bacteria strain (F-) that lacks Cm resistance. In liquid media, we measured conjugation efficiency to be ~1%. When the F+ donor strain was encapsulated within tough hydrogel beads and incubated with recipient bacteria in the surrounding media (2 mL), no transconjugants were detected after 24 hours of co-incubation (LLOD: 1 CFU/2 mL). In additional to conjugation, GMM-derived DNA might reach the environment through diffusion from decayed GMM after cell death. We encapsulated DNA molecules at high concentration (3 × 10^9^ copy/μL) in the beads and measured DNA copy number in the surrounding media using quantitative PCR. There was no DNA molecules leakage as they were perfectly contained (Supplementary Fig. 6) by the alginate-containing DEPCOS, which effectively blocked the diffusion of large biomacromolecules such as DNA^43,44^.

We then investigated the protective effects of the beads on bacterial cells by comparing the resistance of encapsulated cells versus planktonic cells (without bead encapsulation) to a series of chemical and biological stresses (Fig. 2h). Encapsulated bacteria survived to a much greater extent (~350-fold) than planktonic cells in the presence of the aminoglycoside antibiotic kanamycin. Surprisingly, encapsulation also helped cells survive acidic environments (pH 4, ~1700-fold survival improvement). Thus, our robust hydrogels can prevent bacterial conjugation-based HGT and enhance GMM survivability in certain stressful conditions.

To test whether bacterial cells stay metabolically active inside the tough-hydrogel-coated beads, we encapsulated ~30,000/mL cells in each bead and grew them for 24 h (Supplementary Fig. 7). The number of cells in the beads increased by ~10^5^ fold (~16-17 doublings) and reached stationary phase after 12 hours of incubation, corresponding to a doubling time of ~40 minutes. These data indicate that bacterial cells within the beads are metabolically active and able to divide in the alginate core, which is important for GMMs that must carry out active biological functions^45,46^.

The nanoporous structures of the hydrogel shell and alginate core should allow rapid diffusion of small molecules and ions^26,47^ while blocking out large biopolymers such as DNA (Supplementary Fig. 6) and proteins^26^. The anionic nature of alginate in both components further restrict the diffusion of highly charged molecules such as tobramycin^48^ and kanamycin (Fig. 2f). Combining the tough hydrogel shell and the alginate core as a whole system, we observed that mildly charged small molecules can quickly diffuse into the beads (Supplementary Fig. 8). To determine whether encapsulated bacteria can respond to these stimuli, we encapsulated bacteria containing a genetic construct that expresses GFP in response to aTc induction. We then incubated the beads at 37°C in the absence of aTc or in the presence of 200 ng/mL aTc. We found that encapsulated cells exposed to aTc exhibited a 50-fold increase in green fluorescence compared with encapsulated cells not exposed to aTc, which was comparable to the fold-induction seen in liquid cultures (Fig. 3a). Thus, gene expression in cells encapsulated in tough hydrogels can be exogenously controlled by chemical inducers. In addition, significant activation of gene expression could still be observed in ready-to-use beads stored at 4°C for 14 days (Supplementary Fig. 9), which is comparable to current state-of-the-art whole-cell biosensors for field applications^49,50^, such as hydrogel-based^33,51^ and liquid-in-a-cartridge devices^52^. Additionally, to demonstrate the sensing versatility of our system, we showed that a larger and more charged molecule (heme, physiological charge: 3+) with physiological importance could be easily detected using an engineered probiotic *E coli* (Fig. 3b).

**Fig. 3.**
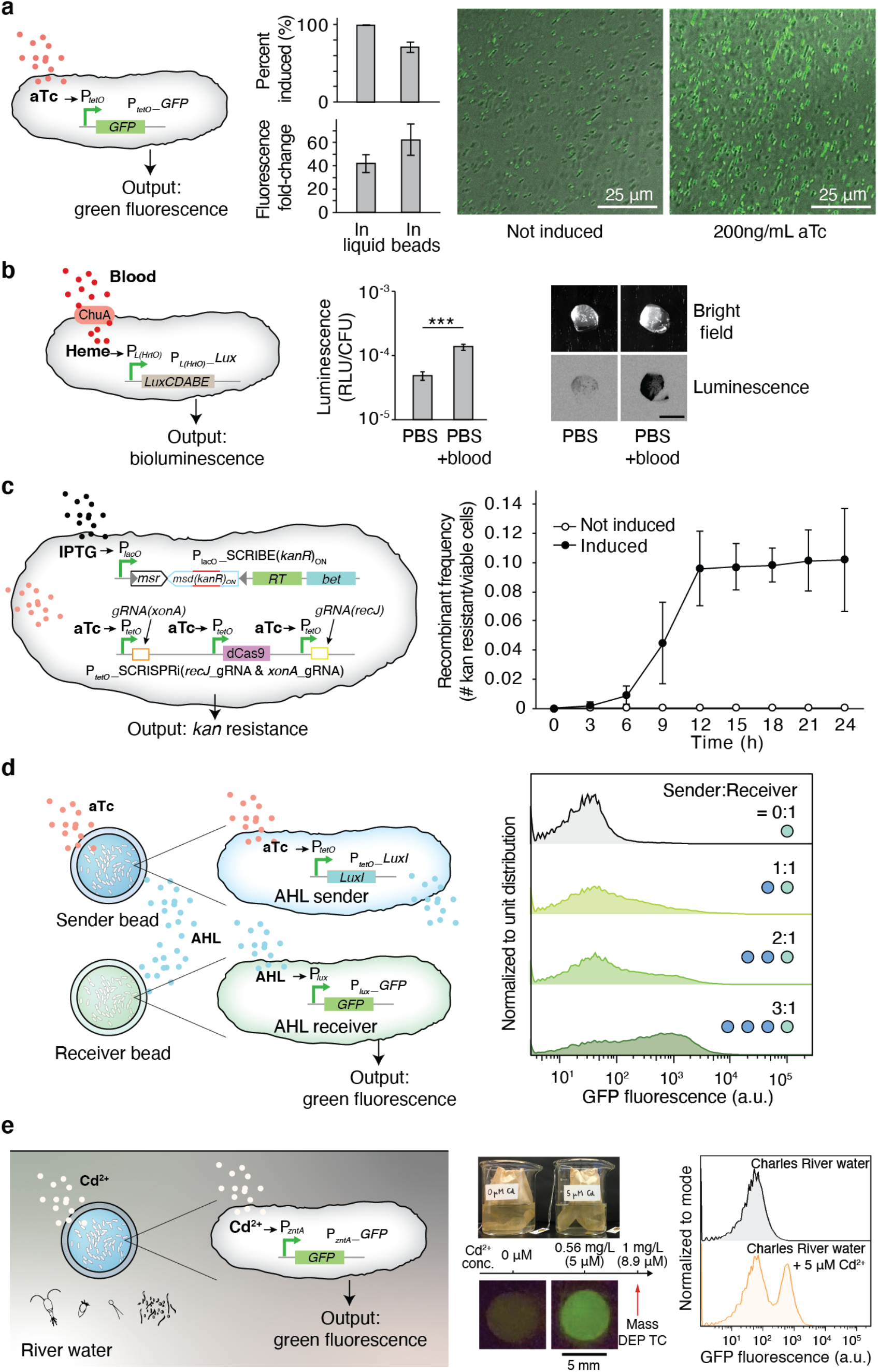
Responses of encapsulated bacterial cells to external stimuli. **(a)** Left: Schematic of GFP expression under the control of an aTc-inducible promoter. Center: Flow cytometry analysis of GFP expression in liquid culture and in hydrogel beads. Data are mean ± s.d. (*n* ≥ 6) Right: Confocal images of beads encapsulating the aTc-sensing *E. coli* strain with and without 200 ng/mL aTc. **(b)** Left: A Heme sensing strain which sense heme and generate bioluminescence as an output^53^. The heme released from blood is transported into the cell by ChuA. Middle: Cells retrieved from beads showed a significant increase in luciferase activity. Right: The resulting bioluminescence can be detected with ultra-high sensitivity from intact beads. **(c)** Left: An improved SCRIBE strain using CRISPRi to knock down cellular exonucleases (*xonA* and *recJ*) for enhanced genome editing efficiency via SCRIBE in DH5αPRO^54^. Right: Recombinant frequencies of beads containing the high-efficiency SCRIBE strain induced for a total of 24 hours with or without aTc and IPTG. Data are mean ± s.d. (*n* ≥ 5). **(d)** Left: An AHL sender strain responds to aTc and produces AHL as an output, which later reaches an AHL receiver strain through diffusion and induces GFP expression. Right: Cells retrieved from receiver beads showed various levels of induction corresponding to different AHL sender bead to AHL receiver bead ratios. The data is representative of three independent experiments and normalized to unit distribution (area under the curve). **(e)** Left: Schematic of GFP expression under the control of a cadmium-inducible promoter. Center: Photograph of the heavy metal sensing experiment setup (top). Tea bags containing five beads each were incubated in beakers containing Charles River water with and without 5 μM CdCl_2_. The Massachusetts Department of Environmental Protection toxic limit for CdCl_2_ is 1 mg/L, corresponding to 8.9 μM. Beads retrieved after 6 hours showed green fluorescence (bottom). Right: Flow cytometry analysis of encapsulated cells responding to cadmium ions in Charles River water (*n* ≥ 3 for all panels). The three flow cytometry panels are each representative of at least four experiments with similar results. Data are normalized to mode (peak value).

We then tested whether bacteria containing a genomically encoded memory system that requires cell division to function would be able to record information within the beads. Recording information on genomic DNA is advantageous in that DNA is a highly stable information storage medium (turnover time up to weeks in aquatic environments and years in soil^55^), information can be retrieved after cell death, and is amenable to multiplexing^56^. We used our SCRIBE platform^45,54^ for targeted *in vivo* genome editing to record information in encapsulated GMMs. The SCRIBE circuit was designed so that IPTG and aTc controlled the expression of Beta recombinase and the CRISPRi system, respectively; in this design, gene editing of the *kanR* gene records chemical exposure (Fig. 3c, left). We exposed beads containing SCRIBE bacteria to IPTG and aTc over 48 hours and found increasing numbers of bacteria acquired kanamycin resistance over the first 12 hours (Fig. 3c, right). The high recombinant frequency (~10%) by 12 hours is comparable to results obtained using liquid cultures of non-encapsulated bacteria^54^, and the plateau in recombination frequency after 12 hours corresponds to growth saturation (Supplementary Fig. 7). This DNA-encoded memory is stable and can be retrieved at the end of the testing period and even after cell death without constant monitoring by electronics.

Communication between GMMs in beads can be used to implement computation with higher complexity, division of labor, and signal integration/amplification^57,58^. To demonstrate this capability, we showed that different *E. coli* strains contained within beads could communicate with each other via quorum-sensing “chemical wires”^59^. An acyl homoserine lactone (AHL) sender strain^60^ and an AHL receiver strain were encapsulated in separate beads and incubated together in 1 mL of Luria-Bertani (LB) media plus carbenicillin (Fig. 3d, left). Upon receiving externally added aTc, the sender bead produced AHL, which induced GFP expression in the neighboring receiver bead. The receiver beads exhibited intensified fluorescence (4-, 12-, and 21-fold-increase for 1, 2, and 3 sender beads, respectively) as more sender beads were used (Fig. 3d right, and Supplementary Fig. 10). These results demonstrate that DEPCOS can enable a modular and distributed strategy for the collective execution of complex tasks based on cell-to-cell communication using multiple beads with different GMMs.

Finally, to demonstrate that encapsulated bacteria can function in a real-world setting, we used an *E. coli* strain to detect the presence of metal ions in water samples from the Charles River, such as cadmium. Cd^2+^ is a well-known and widespread environmental contaminant that can adversely affect human health^61^. Specifically, we used ZntR, a transcriptional regulator activated by metal ions (Zn^2+^, Pb^2+^, Cd^2+^) and activates the promoter P*zntA*, to express GFP^62^. We characterized the induction of P*zntA* by Zn^2+^, Pb^2+^, and Cd^2+^ in liquid cultures of *E. coli* harboring the plasmid pEZ074 (P*zntA*-GFP construct) (Fig. 3e and Supplementary Fig. 11a). While encapsulated in hydrogel beads and incubated in LB media for a total of three hours, cells produced green fluorescence intensities proportional to Cd^2+^ concentrations (Supplementary Fig. 11b).

Next, hydrogel-bacteria beads (pre-soaked in 4x LB) were incubated in water samples extracted from the Charles River having exogenously added Cd^2+^ (Figure 3e, center). The hydrogel-bacteria beads were placed in tea bags to facilitate easy deployment and retrieval. Exposure to 5 μM CdCl_2_ resulted in the emergence of a cell population expressing high levels of GFP (Figure 3e, right), indicating successful detection of cadmium ions. These results were confirmed visually under blue light: beads exposed to 5 μM CdCl_2_ exhibited strong green fluorescence (Fig. 3e, center, and Supplementary Fig. 11c, p < 0.05; Student’s t test). Importantly, the high sensitivity of this system to detect 5 μM CdCl_2_ is relevant to real-world use, as it is below the 8.9 μM (1 mg/L) standard defined by the Massachusetts Department of Environmental Protection as the maximum concentration of cadmium allowed in waste water^32^. Thus, these results highlight the potential of physically biocontained bacteria to detect toxic levels of heavy metals in environmental settings.

To date, the only commercially available GMMs used as environmental sensors are confined in sealed vials into which water samples are manually injected^64^. To enable the environmental deployment of GMMs as biosensors and bioremediation devices, new strategies are needed that allow for interactions with the surrounding environment while maintaining containment of GMMs. Tough hydrogel scaffolds provide a highly hydrated environment that can sustain cell growth, protect cells from external stresses, and allow small molecules to diffuse between the interior and exterior of the device. Although previous work showed the long-term physical containment of bacteria by core-shell hydrogel microparticles, it did not demonstrate biological activity, robust sensing, or high mechanical toughness^32,48,65^. To the best of our knowledge, no reports have demonstrated robust physical containment while still permitting sensing and cell growth, thus overcoming the major limitations for the deployment of GMMs into the real world.

In conclusion, by combining two types of hydrogels into a core-shell structure, we have developed a reliable strategy for the physical containment and protection of microbes that are genetically engineered with heterologous functions. We showed that encapsulated cells could sense environmental and biomedical stimuli, record exogenous signals into genomically encoded memory, and communicate with each other via quorum-sensing molecules. Finally, we showed that heavy-metal-sensing bacteria can be incorporated into our hydrogel beads and successfully detect cadmium ions in Charles River water samples.

We anticipate that the DEPCOS containment platform can enable the deployment of microbes engineered with synthetic gene circuits into real-world scenarios. For example, encapsulated GMMs could be used to detect explosives^66^ or monitor exposure time to toxic chemicals^67^ without potential escape into the wild. In addition, the geometry of DEPCOS could be adapted to meet the design specifications of desired applications, such as wearables^36^. Future work will be focused on automating the manufacturing process to provide precise control over the device size and geometries in order to accommodate various physical environments and improve the miniaturization and scalability of the platform. We will also explore the incorporation of selective diffusion barriers and extreme pH resistance capabilities into the hydrogels to enable encapsulated microbial populations to survive in harsh environments, such as during transit through the human GI tract for detecting disease-relevant biomarkers. Another future challenge lies in devising large-scale standardized tests to determine whether encapsulated organisms can be contained, yet function robustly in harsh real-world scenarios, and not just in simulated laboratory settings.

## Materials and Methods

### Bacterial Strains and Plasmids

A complete listing of bacterial strains and plasmids including their sources can be found in Supplementary Table 1 and Supplementary Table 2. Specifically, pEZ055, pEZ058, and pEZ074 were constructed on a high copy number plasmid (pZE12) backbone carrying a green fluorescence protein (GFP) reporter gene and transformed into DH5αPRO cells. For the aTc-inducible plasmid (pEZ055), the original pZE12 P_LlacO-1_ promoter was substituted by P_LtetO-1_. For the AHL-sensing plasmid (pEZ058), the P_lux_ promoter was PCR amplified and cloned into pZE12 by substituting P_LlacO-1_ promoter via Gibson Assembly. For the heavy-metal-sensing plasmid (pEZ074), the P_zntA_ promoter was PCR amplified from DH5αPRO *E. coli* genomic DNA and cloned into pZE12 by substituting P_LlacO-1_ promoter via Gibson Assembly.

### Manufacturing the Alginate Cores

5 w.t.% alginate solution was made by dissolving medium viscosity alginate (Sigma-Aldrich A2033) in milliQ water followed by autoclaving at 120 °C for 20 minutes to ensure sterility. A fresh bacterial culture (10^8^~10^9^ cells/ml in LB plus antibiotics) was then mixed with the alginate solution in a one to one volume ratio to reach a final alginate concentration of 2.5 w.t.%. This bacteria-alginate premix was loaded into a syringe and disposed onto parafilm to form bead-like droplets with controllable diameters ranging from 2.5 mm to 5 mm. The droplets were solidified by immersing them in 5% w.t. CaCl_2_ (an ionic crosslinker, Sigma-Aldrich 223506) solution for 15 minutes.

### Coating with Tough Hydrogel

A precursor solution composed of 2 w.t.% alginate, 30 w.t.% acrylamide (AAm; Sigma-Aldrich A8887), 0.046 w.t.% ammonium persulphate (APS; Sigma-Aldrich A3678), and 0.015 w.t.% N,N-methylenebisacrylamide (MBAA; Sigma-Aldrich 146072) was thoroughly de-gassed. Before the coating process, the viscous precursor solution was mixed with an accelerator, N,N,N’,N’-tetramethylethylenediamine (TEMED; Sigma-Aldrich T9281; 0.0025 times the weight of AAm) to form a fast-curable pre-gel solution. Alginate cores from the previous section were dipped into the pre-gel solution to form a tunable thin shell layer of 100~1000 microns surrounding the core under a nitrogen atmosphere. To stabilize the shell layer, the hydrogel then was immersed in a MES buffer (0.1 M MES and 0.5 M NaCl, pH 6.0) together with cross-linkers and catalysts including 0.00125 w.t.% 1-ethyl-3-(3-dimethylaminopropyl)carbodiimide (EDC), 0.000375 w.t.% *N*-hydroxysuccinimide (NHS), 0.00075 w.t.% adipic acid dihydrazide (AAD) to form the covalent bonding between the alginate and polyacrylamide network for 3 hours.

### Retrieval of Bacterial Cells

After experiments described in following sections, beads were retrieved from liquid and the tough shell around the alginate core was carefully removed with razor blade and tweezers. The cores were then placed in tubes containing 1 mL phosphate-buffered solution (PBS, Research Products International) and homogenized with 5 mm stainless steel beads on a Tissue Lyzer II (Qiagen 85300) at 30 Hz for 30 minutes. To quantify cell density, homogenized samples were serially diluted (10x) and plated on LB plus antibiotics agar plates. Colony forming units (CFU) were counted after overnight incubation at 37°C.

### Growth of Bacteria in Beads

Beads containing EZ055 were incubated in LB plus carbenicillin at 37°C. At each time point (every 3 hours for a total duration of 24 hours, see Supplementary Fig. 7), cells were retrieved from beads and plated on LB plus carbenicillin agar plates to perform CFU counting.

### Compression Test

The compression of hydrogel beads was carried out using a mechanical testing machine (X N; Zwick/Roell Z2.5). The samples were compressed by two rigid flat substrates at a loading speed of 2 mm min^−1^. As the beads will be immersed in liquid in all practical applications, the mechanical properties (force and displacement) were determined in the swollen state. This was carried out by keeping the bead immerged in LB up until the measurement. The approximate engineering stress is defined as:

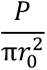

where r_0_, and P are the initial radius of the bead, and the magnitude of the compressive load, respectively^40,68^. After test, beads were incubated in LB overnight and plated to detect potential cell leakage from compression.

### Controlling GRO Life Span

GROs (rEc.β.dC.12’.ΔtY and LspA.Y54β) were grown in LB plus 1 mM p-azido-phenylalanine (pIF, Santa Cruz Biotechnology SA-289923), 0.02% L-arabinose (L-ara), and carbenicillin at 30°C overnight^21^, washed twice with PBS to remove pIF and L-ara, and encapsulated in hydrogel beads. GRO beads were then incubated in LB plus carbenicillin with or without 1 mM pIF at 4°C overnight to allow pIF infusion. At t = 0, beads were placed in 50 mL LB medium and incubated at 30°C for 12 h, 24 h, and 48 h. Cells were retrieved at given time points and plated on LB plus carbenicillin with 1 mM pIF and 0.02% L-ara agar plates for CFU counting. Survival rates were calculated by normalizing CFU counts to t = 0.

### Detecting GRO Escape

GROs (rEc.β.dC.12’.ΔtY and LspA.Y54β) were grown in LB plus 1 mM p-azido-phenylalanine (pIF), 0.02% L-arabinose (L-ara), and carbenicillin at 30°C overnight, washed twice with PBS to remove pIF and L-ara, and encapsulated in alginate beads with or without tough hydrogel coating. The beads were then incubated in 5 mL LB plus carbenicillin at 30°C with 200 rpm shaking for 3 days. Media from each tube was plated on LB plus carbenicillin plates for CFU counts.

### Environmental Insult Experiments

For antibiotics and acidic condition treatments, beads containing EZ074 cells were incubated in 1 mL of LB plus carbenicillin at 37°C for 12 hours as shown in Supplementary Fig. 12. This step acts as an outgrowth phase to bring cell densities in the different beads to a similar level (~10^8^ per bead). At t = 0, culture media was switched to LB plus 30 μg/ml kanamycin and LB at pH 4, respectively. At the end of the experiments, beads were retrieved from liquid media and cells were harvested for CFU counting.

### Bacterial Conjugation

The F’ plasmid (containing chloramphenicol resistance) donor strain CJ236 was encapsulated in beads and underwent overnight outgrowth in LB without antibiotics. The donor beads were placed in 2 mL of LB and co-cultured with recipient strain rcF453 (with streptomycin resistance). After 24 hours of incubation at 37°C (shaking at 100 rpm), the surrounding media was plated on LB plus streptomycin (Sm, 25 μg/mL) and LB plus streptomycin (25 μg/mL) and chloramphenicol (Cm, 12.5 μg/mL). The conjugation efficiency was calculated as:

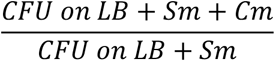

### DNA Escape Test

A linear DNA fragment encoding a GFP transcriptional unit (~1k bp) was amplified using PCR and encapsulated in the hydrogel beads at 3e9 copy/μL. Soluble DNA in the surrounding media after 72-hr incubation was quantified using qPCR (Roche LightCycler 96) with an optimized amplicon (314 bp). A standard curve was constructed using a serial dilution (10x) of the same fragment.

### Small Molecule Diffusion Assay

Two fluorescent dyes, rhodamine B and fluorescein, were used as surrogates for small molecule diffusion assay. Coated beads were soaked in dye solutions for various time periods, weighted, and transferred into 2 mL of PBS and incubated in dark for 24 hr. The fluorescence of the PBS at equilibrium was measured (494/521 nm and 540/625 nm with a Synergy H1 Hybrid Multi-Mode Reader, BioTek Instruments), calibrated by the total weight of PBS plus bead, and normalized to the saturated maximum incubation period in dyes (24 hr).

### aTc Induction in Beads

Beads containing EZ055 were incubated in LB plus carbenicillin and 200 ng/mL aTc at 37°C for 8 hours. The bead was then retrieved and sliced with a sharp razor blade at thickness of ~0.5 mm. The sliced sample was then imaged with a Zeiss LSM 700 confocal microscope with excitation wavelength at 488 nm and emission wavelength at 515 nm. To test inducibility after long-term storage, beads were kept at 4°C over the course of 30 days/ At each time point, beads were retrieved and induced in LB plus carbenicillin and 200 ng/mL aTc at 37°C for 8 hours. Fluorescence profiles were characterized using a Synergy H1 Hybrid Multi-Mode Reader (488 nm excitation, 530/30 detection, BioTek Instruments).

### Heme Sensing in Beads

Defibrinated horse blood (Hemostat Laboratories DHB030) was used as the source of blood and was lysed by first diluting 1:10 in simulated gastric fluid (SGF) (0.2% NaCl, 0.32% pepsin, 84 mM HCl, pH 1.2) to release heme. This stock solution was diluted to 300 ppm in PBS right before experiments. Beads were placed in PBS or PBS + blood for 8 hours at 37°C. Cells were retrieved and measured for luminescence using a Synergy H1 plate reader. The relative luminescence units were normalized by CFU (measured through plating). Luminescence images of intact beads were acquired using a ChemicDoc Touch Imageing System (Bio-Rad).

### Memory of Chemical Exposure (SCRIBE) in Beads

An higher efficiency version of SCRIBE (Synthetic Cellular Recorders Integrating Biological Events) was used in this study^45,54^. The SCRIBE strain was encapsulated in tough hydrogel beads and incubated in LB media with carbenicillin (100 μg/mL), chloramphenicol (25 μg/mL), aTc (100 ng/ml), and IPTG (1 mM) at 37°C. A control group was incubated using the same conditions but without the inducers (aTc and IPTG). At given time points, cells were retrieved from the beads and plated on LB plus kanamycin (50 μg/mL) agar plates as well as LB plus carbenicillin and chloramphenicol agar plates. The recombinant frequency was calculated by dividing the colony count on the LB plus kanamycin plate (*kan*-resistant cells) by the colony count on the LB plus carbenicillin and chloramphenicol plate (total viable cells).

### Quorum Sensing between Beads

Beads containing the AHL sender strain (AYC261) and AHL receiver strain (EZ058) were placed in 1 mL LB plus 250 ng/ml aTc in a 12-well plate at specific ratios (sender:receiver = 0:1, 1:1, 2:1, and 3:1). After 24 hours of incubation at 37°C, we retrieved and diluted AHL receiver cells 1:20 into phosphate-buffered solution (PBS, Research Products International) and ran them on a BD-FACS LSRFortessa-HTS cell analyzer (BD Biosciences). We measured at least 20,000 cells for each sample and consistently gated by forward scatter and side scatter for all cells in an experiment. GFP intensity was measured on the FITC channel (488-nm excitation laser, 530/30 detection filter). Data from flow cytometry is normalized to unit distribution (normalized to area under the curve).

### Heavy Metal Sensing

To test inducibility of the Zn/Pb/Cd sensing strain (EZ074) in liquid, cells were grown overnight and diluted 200x in fresh LB plus 300 μM ZnCl_2_, 100 μM Pb(NO_3_)_2_, and 10 μM CdCl_2_, respectively on a 96-well plate. After 3 hours of incubation at 37°C, we retrieved bacterial cells and analyzed their GFP profile with flow cytometry. For testing inducibility in beads, EZ074 was encapsulated in tough hydrogel bead and incubated overnight in LB media with carbenicillin at 4°C. Before the experiment, beads were incubated at 37°C for 12 hours for bacterial cell outgrowth. Hydrogel beads were then placed in fresh LB medium with carbenicillin and corresponding metal ions at given concentrations and incubated at 37°C for 3 hours. Bacterial cells were retrieved and analyzed with flow cytometry. Data from flow cytometry is normalized to mode (normalized to peak value), which allows the visualization of differences in relative percentages of cell populations of interest.

### Metal Sensing in Charles River Water

Beads containing EZ074 were incubated in 4x LB media at 4°C overnight to reach equilibrium. At t = 0, beads were placed in teabags and transferred to beakers containing 100 mL of fresh Charles River water with or without 5 mM CdCl_2_. After 6 hours of incubation at room temperature, cells were retrieved and analyzed with flow cytometry. Data from flow cytometry is normalized to mode.

## Acknowledgments

The authors thank Dr. Fahim Farzadfard for proving the SCRIBE strains and Dr. Mark Mimee for providing the heme sensing strain. We thank Shaoting Lin, Dr. Cesar de la Fuente-Nunez, Dr. Nathaniel Roquet, Dr. Robert Citorik, and Dr. Sebastien Lemire for useful discussions. This work was supported by the NIH New Innovator Award (1DP2OD008435), NIH National Centers for Systems Biology (1P50GM098792), the U.S. Office of Naval Research (N000141310424), National Science Foundation (EFMA-1935291, CMMI-1661627), Office of Naval Research (N00014-17-1-2920), and the U.S. Army Research Office through the Institute for Soldier Nanotechnologies at MIT (W911NF-13-D-0001), the J-WAFS Graduate Student Fellowship (T.-C.T.), and the Defense Advanced Research Projects Agency.

## Author contribution

T.-C.T., E.T., X.L., X.Z., and T.K.L. conceived and designed the research; T.-C.T., E.T., X.L., and H.Y. performed encapsulation and mechanical testing experiments; T.-C.T. and E.T. performed genetic circuit experiments; T.-C.T., E.T. and A.J.R. performed GRO experiments; T.-C.T. and E.T. performed river water experiments; T.-C.T., E.T., X.L., K.Y., A.J.R., F.J.I., X.Z., and T.K.L analyzed the data and wrote the manuscript.

