## Supplementary Materials for "Tough Hydrogel-Based Biocontainment of Engineered Organisms for Continuous, Self-Powered Sensing and Computation"

Supplementary Tables & Figures

Supplementary Table 1 | List of bacterial strains used in this study

| Name | Strain code | Construction method | Genotype | Used in |
|---|---|---|---|---|
| GRO, pIF auxotroph strain | rEc.β.dC.12'.ΔtY | Ref[1] | MG1655 | Fig. 2 |
| GRO, pIF auxotroph strain | LspA.Y54β | Ref[1] | MG1655 | Fig. 2 |
| aTc sensing strain | EZ055 | DH5αPRO cells transformed with the pEZ055 plasmid | DH5αPRO | Fig. 2<br>Fig. 3 |
| F' plasmid donor strain | CJ236 | Acquired from NEB | K12 | Fig. 2 |
| F' plasmid recipient strain | rcF453 | Spontaneous resistant mutants generated from plating and re-streaking MG1655 on LB+Sm plate | MG1655 | Fig. 2 |
| Zn/Pb/Cd sensing strain | EZ074 | DH5αPRO cells transformed with the pEZ074 plasmid | DH5αPRO | Fig. 2<br>Fig. 3 |
| SCRIBE *kanR*$_{OFF}$ reporter strain | F144 | Ref[2] | DH5αPRO *galK::kanR*$_{W28TAA,\ A29TAG}$ | Fig. 3 |
| AHL sender strain | AYC261 | Ref[3] | DH5αPRO | Fig. 3 |
| AHL receiver strain | EZ058 | DH5αPRO cells transformed with the EZ058 plasmid | DH5αPRO | Fig. 3 |
| Heme sensing strain | mm1560 | Ref[4] | Nissle 1917 | Fig. 3 |

Supplementary Table 2 | List of plasmids used in this study

| Name | Plasmid code | Construction method | Used in |
|---|---|---|---|
| $P_{LtetO-1}$_*gfp* | pEZ055 | See Materials and Methods | Fig. 2<br>Fig. 3 |
| $P_{zntA}$_*gfp* | pEZ074 | See Materials and Methods | Fig. 2<br>Fig. 3 |
| $P_{lacO}$_SCRIBE(*kanR*)on | F944 | Ref[2] | Fig. 3 |
| $P_{TetO}$_CRISPRi(*recJ*_gRNA & *xonA* gRNA) | F1156 | Ref[5] | Fig. 3 |
| $P_{TetO}$_*LuxI* | AYC261 | Ref[3] | Fig. 3 |

Supplementary Fig. 1

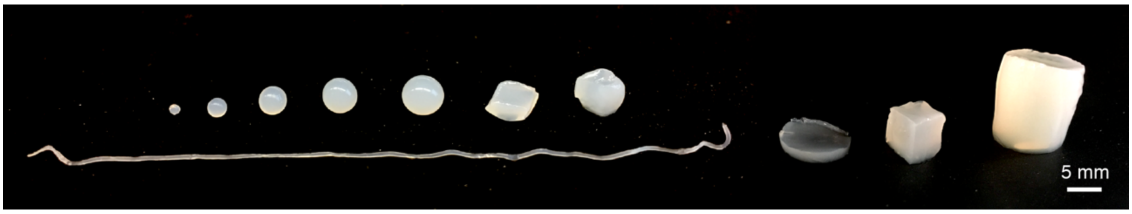


**Supplementary Fig. 1 | Alginate cores in various geometries.** The alginate core used to encapsulate cells can be shaped into spheres with different radii through extrusion with syringes and needles on parafilm followed by crosslinking in calcium chloride solution. Alginate thread was produced by direct extrusion in calcium chloride solution. Disk, cube, and cylinder-like structures can be achieved through cutting.

Supplementary Fig. 2

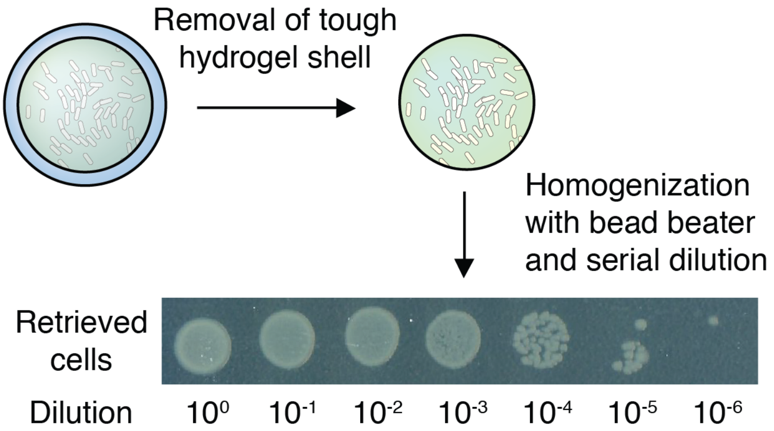


**Supplementary Fig. 2 | Retrieving encapsulated cells.** Retrieval of live cells from the beads immediately following the cross-linking step. Retrieval was performed through removal of the tough shell followed by homogenization and showed recovery levels around 20% (number of retrieved cells/number of encapsulated cells).

Supplementary Fig. 3

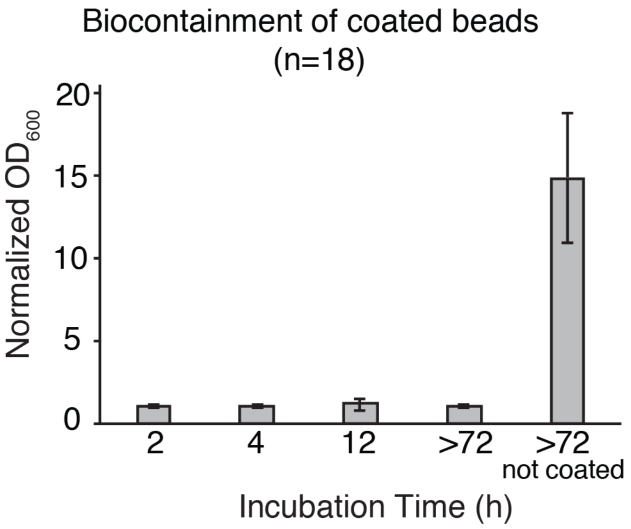


**Supplementary Fig. 3 | Long-term physical containment.** Optical density at 600 nm ($OD_{600}$) measurement demonstrating that the media surrounding coated beads showed no bacterial growth after 72 hours. Data are mean ± s.d. (*n* = 18).

Supplementary Fig. 4

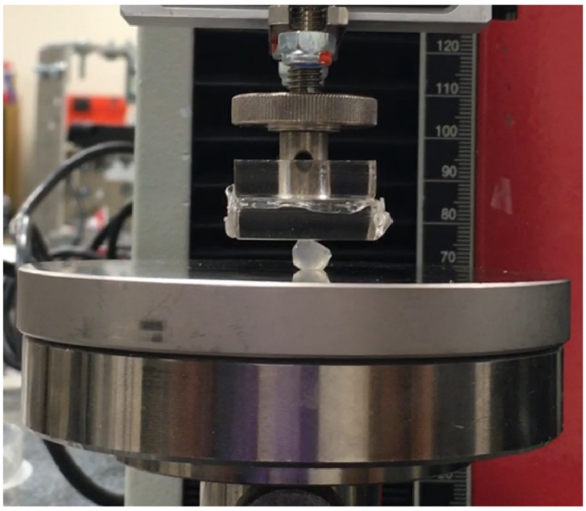

**Supplementary Fig. 4 | Compression test setup with Zwick mechanical tester.** A fully-hydrated hydrogel bead (r = 4 mm) was placed between sterile surfaces and submitted to compressions.

Supplementary Fig. 5

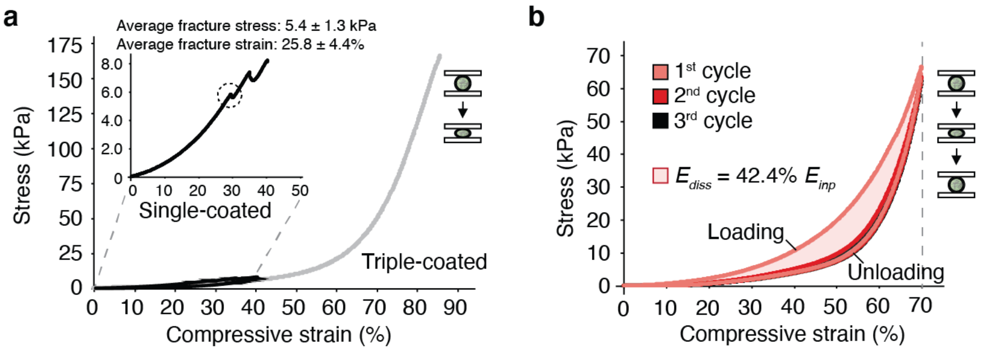


**Supplementary Fig. 5 | Effective stress-strain profiles of the hydrogel beads under compression. (a)** Effective stress-strain curves of single- and triple-coated beads. (**b**) Effective stress-strain curves of cyclic compression of triple-coated beads Effective stress-strain curves were converted from force-displacement curves using the initial dimensions of the beads before compressions[6,7].

Supplementary Fig. 6

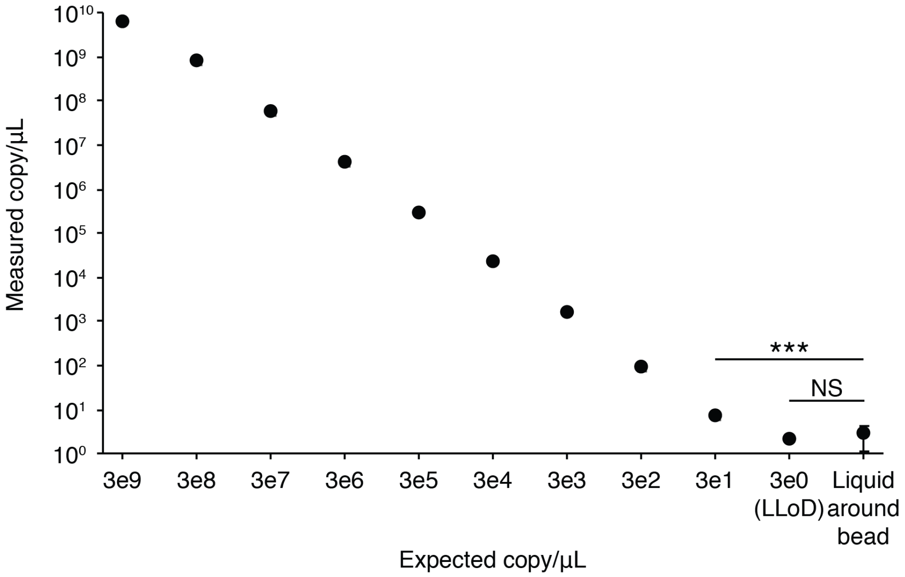


**Supplementary Fig. 6 | DNA containment inside the hydrogel beads.** Linear DNA fragments (977 bp) were PCR-amplified and encapsulated in the hydrogel beads at 3e9 copy/µL. Soluble DNA in the surrounding media after 72-hr incubation was quantified using qPCR. Standards were prepared by serial dilutions. Data are mean ± s.d. (*n* = 3, ***p < 0.001). Lower limit of detection (LLOD) = 3 copy/µL. NS = not significant.

Supplementary Fig. 7

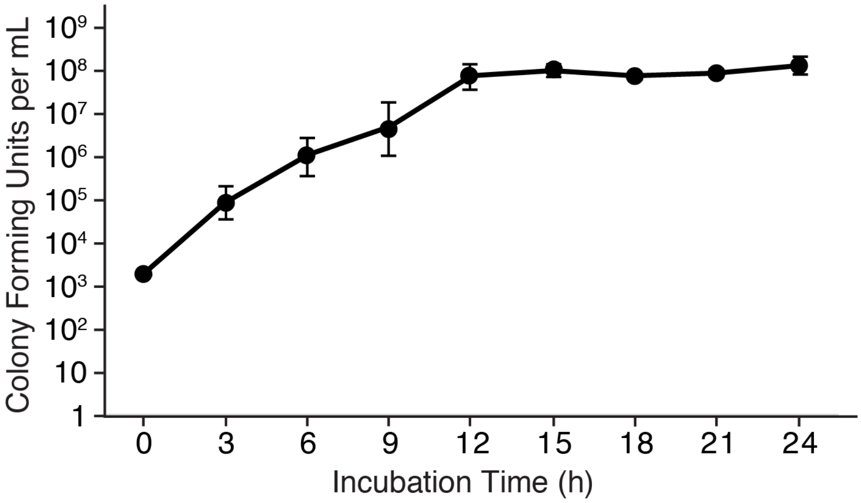


**Supplementary Fig. 7 | Growth curve of bacteria in hydrogel beads.** Cell encapsulated in beads were incubated in LB medium and retrieved at given time points to measure growth over 24 hours. Data are mean ± s.d. ($n$ = 3).

Supplementary Fig. 8

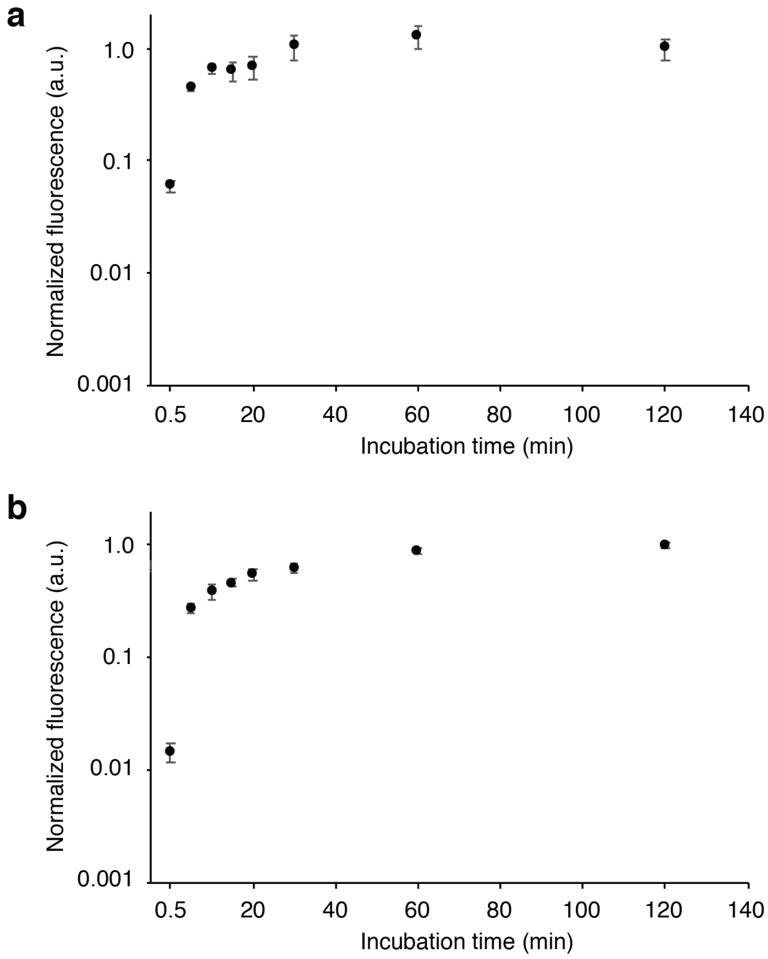


**Supplementary Fig. 8 | Diffusion of small molecules into the hydrogel beads. (a)** Diffusion of a positively charged dye, rhodamine, into the hydrogel beads over a course of two hours. **(b)** The diffusion profile of a negatively charged dye, fluorescein. Data are mean ± s.d. (*n* = 3).

Supplementary Fig. 9

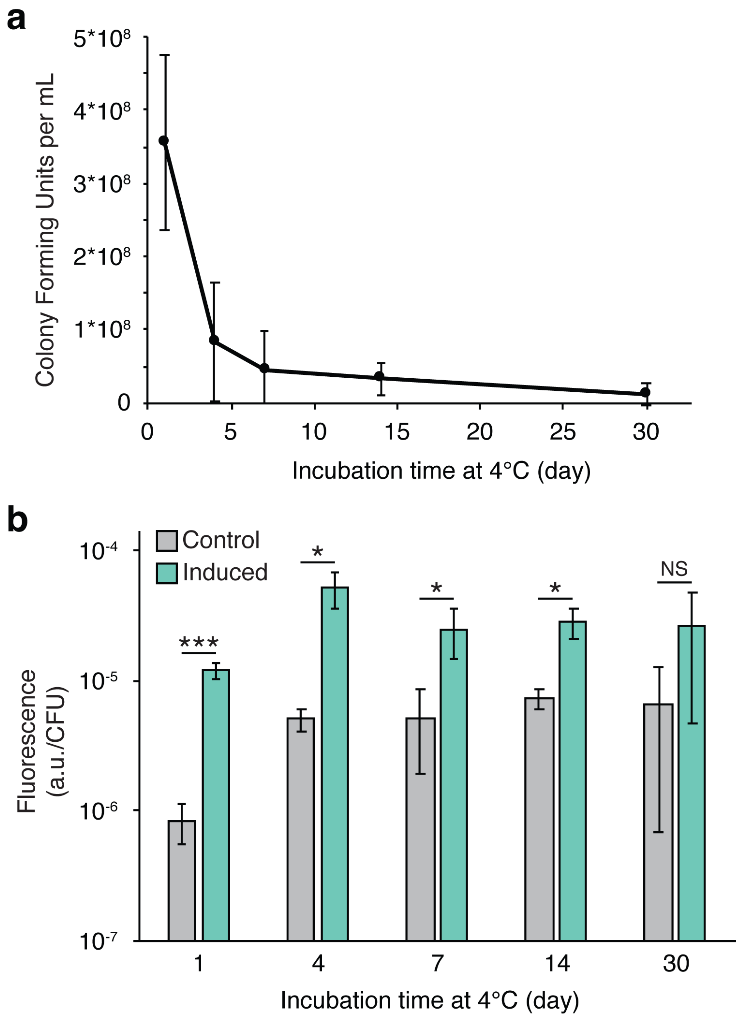


**Supplementary Fig. 9 | Cell survival and inducibility after storage at low temperature. (a)** CFU counts for cells retrieved from hydrogel beads after storage in a refrigerator (4°C) across 30 days **(B)** Comparison of aTc-induced fluorescence profiles of retrieved cells after storage at 4°C for various time periods. Data are mean ± s.d. ($n$ = 3, ***p < 0.001, *p < 0.05, NS: not significant).

Supplementary Fig. 10

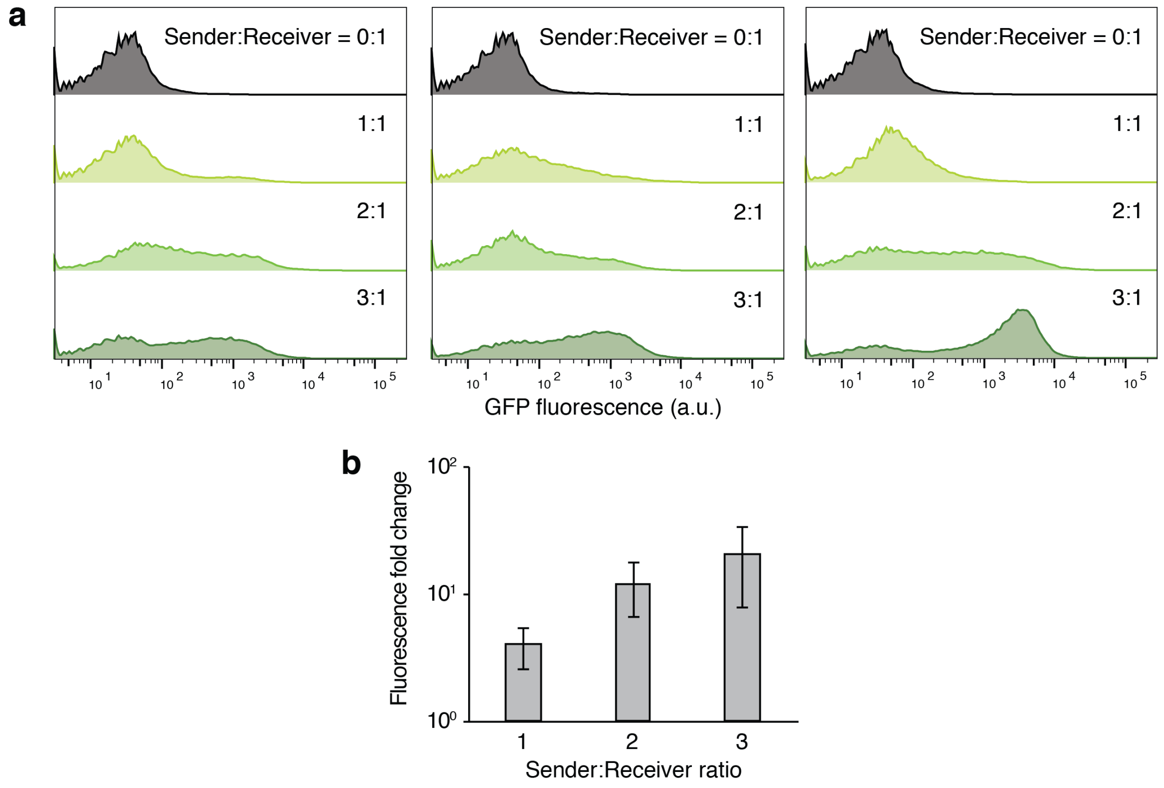


**Supplementary Fig. 10 | Induction of AHL receiver beads by AHL sender beads. (a)** Flow cytometry data of cells retrieved from receiver beads showed various levels of induction corresponding to different AHL sender bead to AHL receiver bead ratios (normalized to unit distribution, three biological replicates). **(b)** Fluorescence fold change of the receiver beads. Data are mean ± s.d. ($n$ = 3).

Supplementary Fig. 11

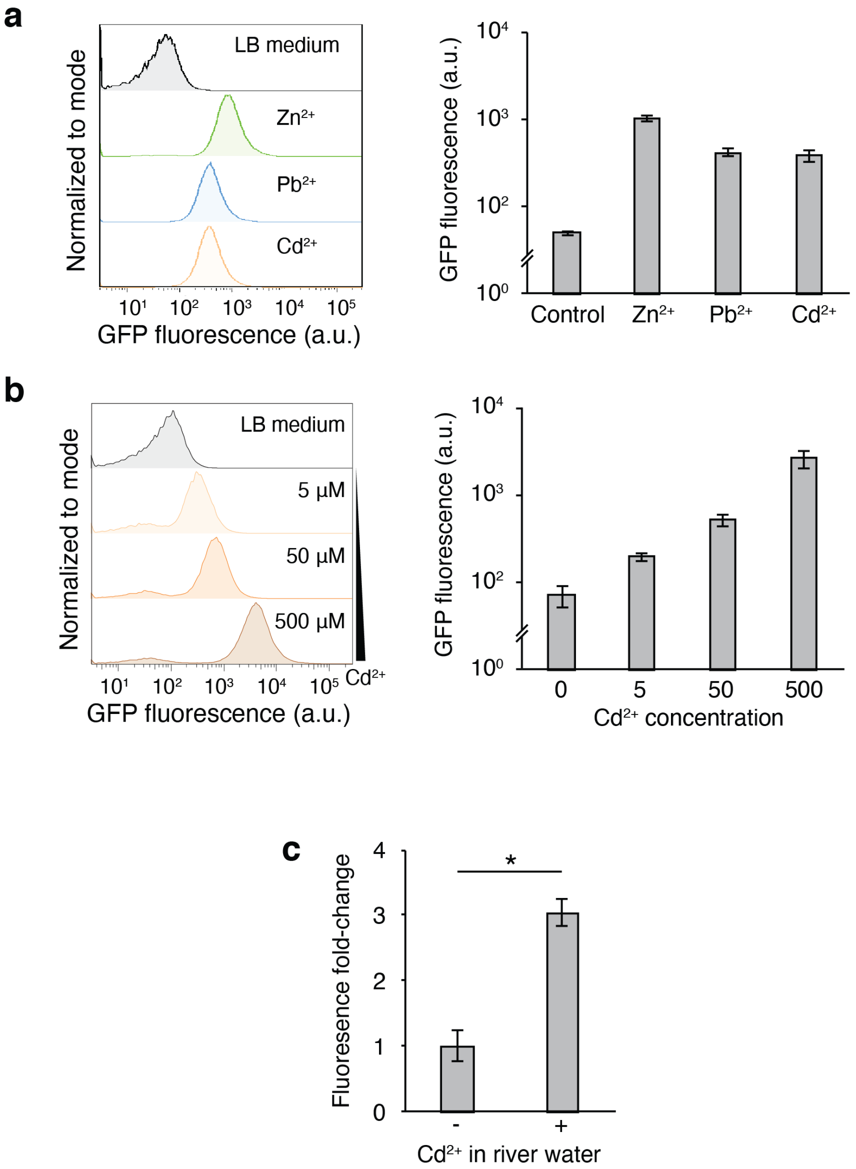


**Supplementary Fig. 11 | Heavy metal sensing in Charles River water samples. (a)** Left: Flow cytometry analysis of the heavy-metal-sensing strain. Bacteria in liquid were exposed for 3 hours to 300 µM $ZnCl_2$, 100 µM $Pb(NO_3)_2$, and 10 µM $CdCl_2$ in LB media, respectively. Right: Mean GFP fluorescence of the heavy-metal-sensing strain. **(b)** Left: Response of the heavy-metal-sensing strain encapsulated in the tough hydrogel capsule to 0 µM, 5 µM, 50 µM, and 500 µM $CdCl_2$ after 3 hours of incubation. Right: Mean GFP fluorescence of the heavy-metal-sensing strain encapsulated in the tough hydrogel capsule. **(c)** GFP fluorescence fold-change of encapsulated cells responding to cadmium ions in Charles River water ($n \geq 3$ for all panels, *$p < 0.05$).

Supplementary Fig. 12

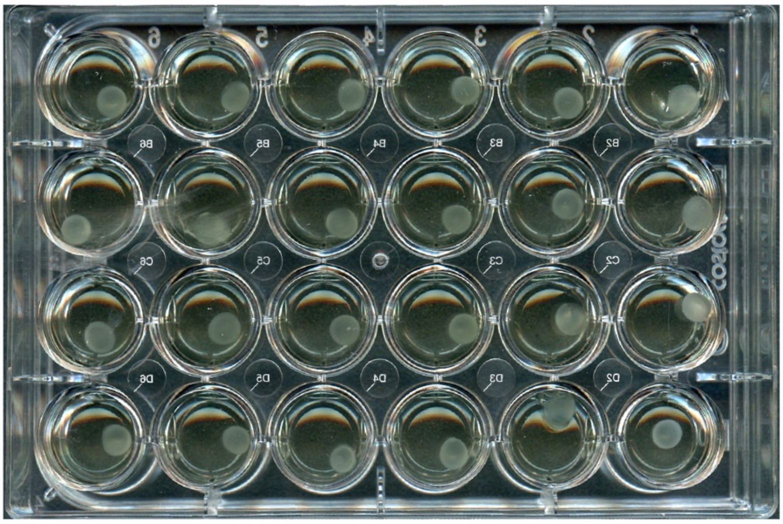

**Supplementary Fig. 12 | The setup of induction and insult experiments.** Each bead containing engineered bacterial cells was incubated in LB + antibiotics in a 24-well plate. After showing no signs of bacterial leakage/escape after overnight incubation, beads were transferred to a new plate and fresh media/chemicals were added to the wells to start experiments.
